# Comparing infiltration rates in soils managed with conventional and alternative farming methods: a meta-analysis

**DOI:** 10.1101/603696

**Authors:** Andrea D. Basche, Marcia S. DeLonge

**Affiliations:** Department of Agronomy and Horticulture, University of Nebraska-Lincoln, 1875 N. 38^th^ Street, Plant Sciences Hall, Lincoln, NE 68503, U.S.A.; Food & Environment Program, Union of Concerned Scientists, 500 12^th^ Street, Suite 340, Oakland, CA, 94607-4087, U.S.A.

## Abstract

Identifying agricultural practices that enhance water cycling is critical, particularly with increased rainfall variability and greater risks of droughts and floods. Soil infiltration rates offer useful insights to water cycling in farming systems because they affect both yields (through soil water availability) and other ecosystem outcomes (such as pollution and flooding from runoff). For example, conventional agricultural practices that leave soils bare and vulnerable to degradation are believed to limit the capacity of soils to quickly absorb and retain water needed for crop growth. Further, it is widely assumed that farming methods such as no-till and cover crops can improve infiltration rates. Despite interest in the impacts of agricultural practices on infiltration rates, this effect has not been systematically quantified across a range of practices. To evaluate how conventional practices affect infiltration rates relative to select alternative practices (no-till, cover crops, crop rotation, introducing perennials, crop and livestock systems), we performed a meta-analysis that included 89 studies with field trials comparing at least one such alternative practice to conventional management. We found that introducing perennials (grasses, agroforestry, managed forestry) or cover crops led to the largest increases in infiltration rates (mean responses of 59.2 ± 20.9% and 34.8 ± 7.7%, respectively). Also, although the overall effect of no-till was non-significant (5.7 ± 9.7%), the practice led to increases in wetter climates and when combined with residue retention. The effect of crop rotation on infiltration rate was non-significant (18.5 ± 13.2%), and studies evaluating impacts of grazing on croplands indicated that this practice reduced infiltration rates (−21.3 ± 14.9%). Findings suggest that practices promoting ground cover and continuous roots, both of which improve soil structure, were most effective at increasing infiltration rates.

## Introduction

There is a need to develop more resilient, multifunctional agricultural systems, particularly given risks posed by climate change to farm productivity and environmental outcomes^1,2,3^. Specifically, water-related risks from increased rainfall variability include soil erosion and water pollution, degradation of soil quality, and reductions to crop yields^4,5,6^. Although soils are vulnerable to water-related risks, they are also being recognized as a medium to mitigate such risk when managed to deliver a wide range of ecosystem benefits, beyond maximizing crop production^7,8^. Thus, designing agricultural systems that improve soils and soil water cycling is one strategy that could help reduce negative impacts of increasing rainfall variability^9,10,11,12^. To this point, global modeling analyses indicate that enhancing soil water storage at a large scale can benefit crop productivity and improve ecosystem services, such as by reducing runoff^13,14^. However, there is a need to identify how to secure such outcomes on the farm-scale, particularly across a range of management practices, environments, and climates.

Emerging interest in how soils can support climate adaptation has increased the urgency to understand the potential benefits of farms shifting from conventional to alternative agricultural practices. Presently, conventional cropping systems typically feature annual crops, leave the soil bare when a cash crop is not growing, have limited crop diversity, and include regular soil disturbance through tillage: within the United States, only approximately 3% of cropland acres are growing a cover crop and 25% are utilizing no-till practices.^15,16,17^ Soil disturbance, a lack of soil cover and limited plant diversity can degrade soils, reducing their ability to withstand rainfall variability through affects such as disrupting aggregation, increasing bulk density, and limiting water holding capacity.^18^ In contrast, management practices such as no-till and cover crops may counter such degradation but remain in the minority.^19^ The limited adoption rates may be in part related to the fact that, in spite of decades of agronomic research surrounding such practices, we are only beginning to understand their potential value for improving key functions related to soil health and water cycling.^18^

A growing body of research suggests that a range of alternative farming practices can contribute to biological, physical and chemical transformations in soil that in turn can increase water storage, improving resilience to droughts, floods, and extreme weather conditions^20,21^. For example, studies have shown that no-till, cover crops and crop rotations can in some cases improve soil carbon content, soil biological activity, and soil physical properties associated with water storage^22-27^. There is also evidence that practices such as introducing perennials and designing diversified landscapes, such as through integrating crop and livestock practices, can improve soils in similar ways, likely by protecting soils and including living roots throughout the year^28-31^. However, because there are a number of different soil water measurements, the effects of specific practices on soil water properties have not previously been well summarized quantitatively^20^.

The primary goal of this analysis was to synthesize published field-experiments investigating impacts of agricultural practices on water infiltration and to gain insight into mechanisms impacting infiltration rates. We focused on soil infiltration rates because, in addition to being critical to mitigating drought and flood risk^29^, they are frequently measured in field experiments and are sensitive to changes in management. Infiltration rates are also closely related to other important characteristics of soils, including physical aspects such as aggregate stability, bulk density, plant available water, as well as chemical and biological aspects including soil carbon, and microbial biomass^20,26,27^. In this study, we considered a range of specific alternative practices that can be adopted on farms, including no-till, cover crops, crop rotations, introducing perennials, and livestock grazing on croplands, compared to more conventional controls (experiments with tillage, no cover crops, monocropping, annual crops, and no grazing). We secondarily explored patterns of additional environmental and management factors (e.g. soil texture, climate indices, and the length of the experiment) that could be modulating observed effects.

## Methods

### Study criteria

We evaluated the effects of various alternative farming practices that can be adopted in otherwise conventional farming systems,^32-34^. We considered zero tillage (*no-till*) as compared to conventional tillage, cover cropping or green manure practices that keep soils covered compared to leaving them bare (*cover crops*), diversified farming (crop rotations, intercropping) as compared to monoculture cropping (*crop rotations*), agricultural systems with mainly perennial compared to annual crop systems (*perennials*), and grazing of croplands versus conventionally harvested or hayed fields (*crop and livestock*) (Figs 1-2 and Table 1). The main criteria for inclusion were field experiments that: 1. Measured and reported steady-state infiltration or the volume of water entering the soil over a designated period; 2. Compared one of the alternative practices of interest relative to select conventional controls in a standardized way.

**Table 1.**
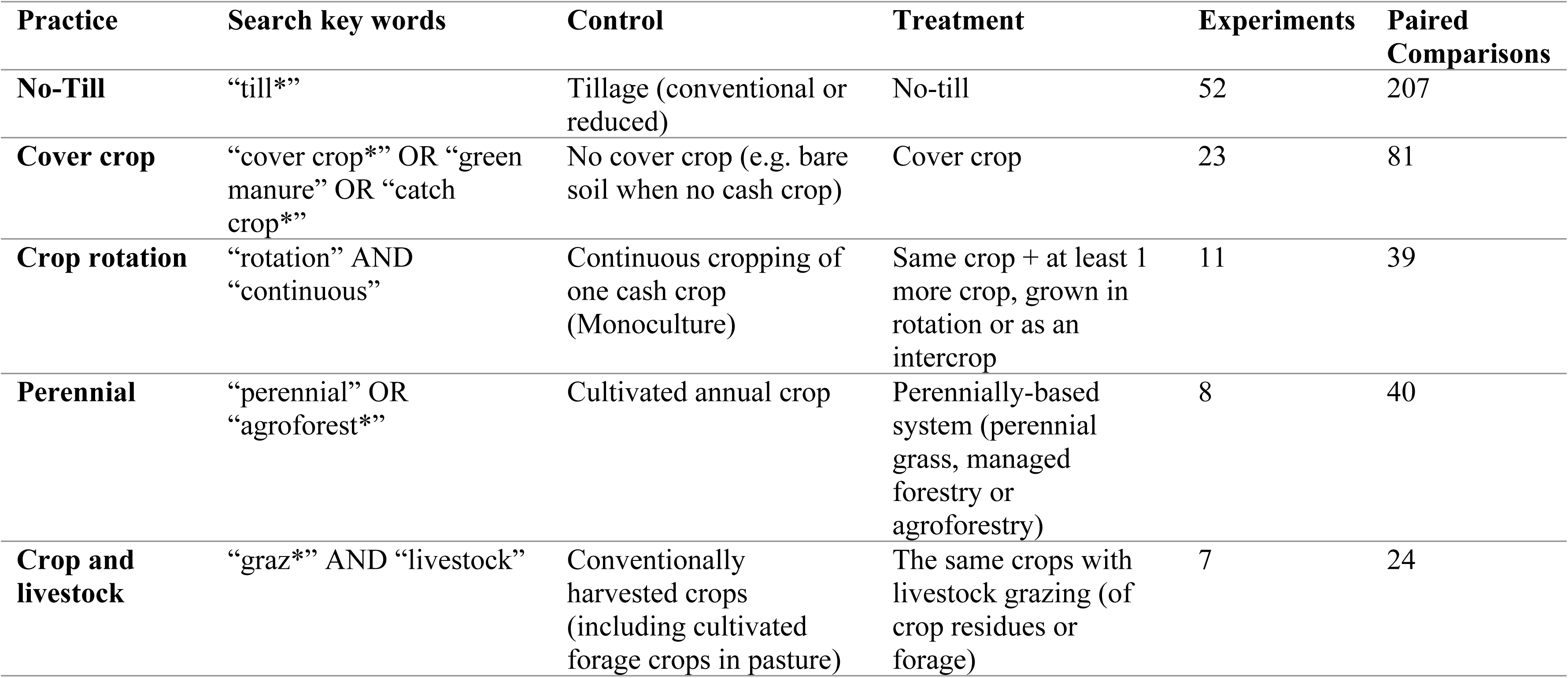
Criteria and results for literature searches for specific agricultural practice comparisons.

**Fig 1.**
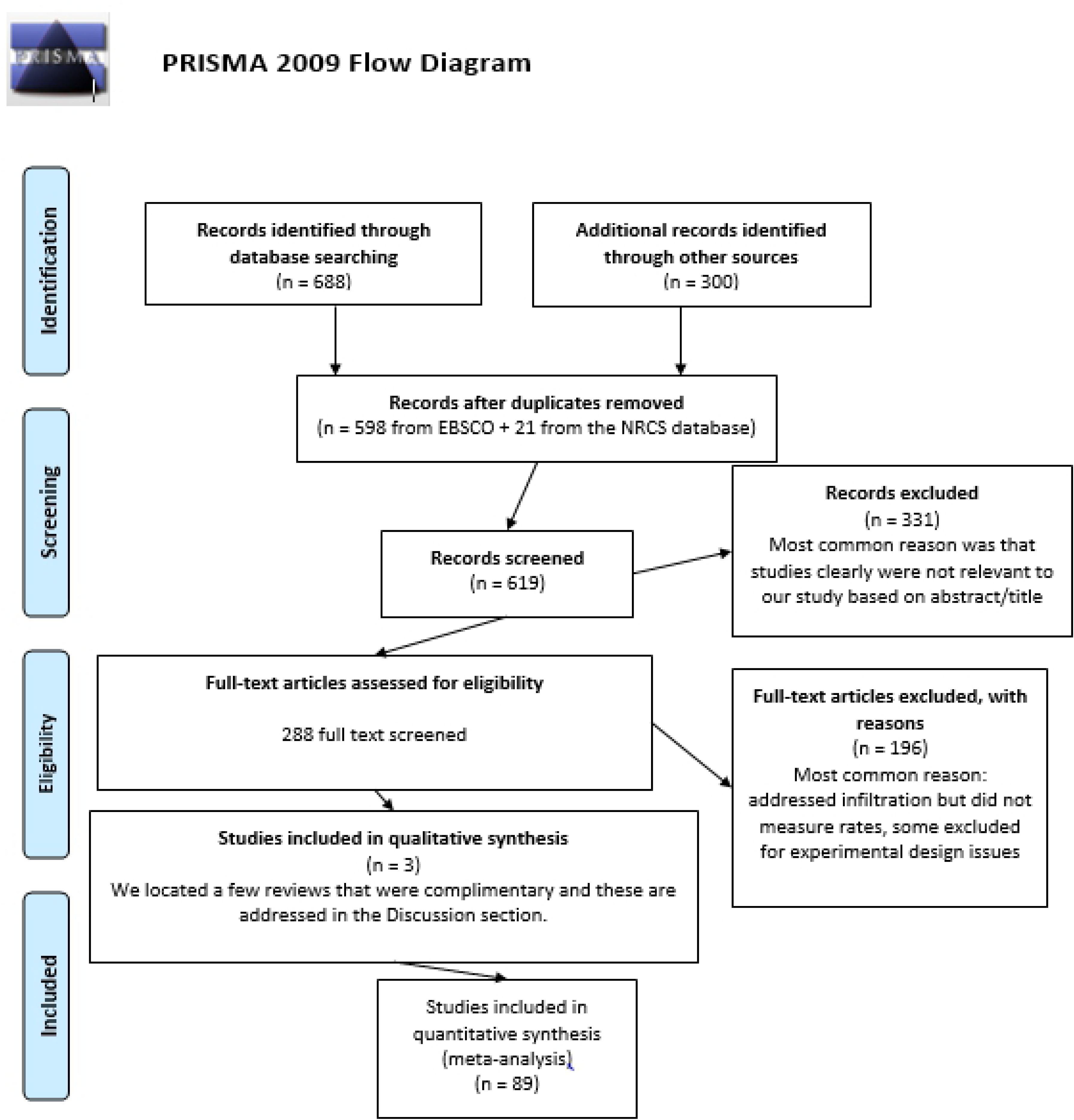
Preferred Reporting Items for Systematic Reviews and Meta-Analysis (PRISMA) Flow Chart describing the search protocol utilized to identify and select published research for this analysis.

**Fig 2.**
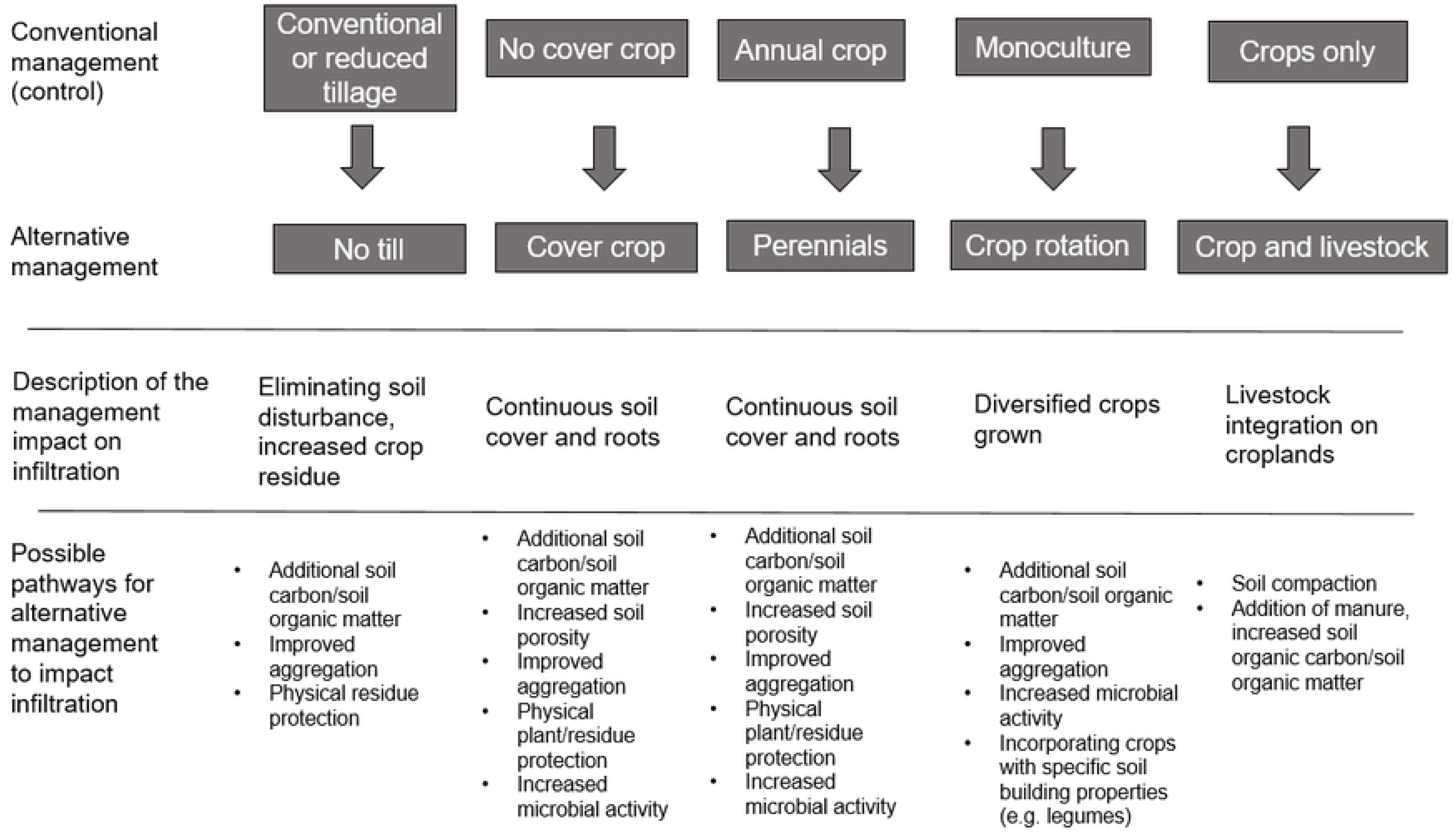
Alternative agricultural practices evaluated in this analysis and the conventional practice controls. The five different practices could alter infiltration through a range of physical, chemical or biological processes, as described

Shown here are the control and treatment conditions for all practice comparisons considered for this study, as well as the number of experiments and specific paired comparisons (response ratios) that met the criteria for inclusion into the meta-analysis. Additional details for each experiment are in Supporting Information (Table 1 in S1 File).

### Literature search

The literature search was conducted using *EBSCO Discovery Service*™ (detailed in Basche and DeLonge^25^) and only included field experiments in English language peer-reviewed literature through 2015 (the earliest publication that met our criteria was from 1978). Keyword strings included “infiltration W1 rate” AND “crop*” for all searches, and additional keywords were used for individual practices (Table 1). These searches returned approximately 700 studies, of which 79 fit our criteria. We used the USDA-NRCS Soil Health Literature database^35^ to find additional papers, leading to 10 more studies for a total of 89 (Table 1). For additional details, see the Supporting Information.

### Management practices

Experiments within each practice were systematically included in the database only if they fit the below additional criteria.

#### No till

Papers identified from the additional search term “till*” were included if experiments clearly included a no-till treatment. We compared any tillage practices – reduced tillage as well as more physically disruptive tillage practices that are typically described as conventional tillage – to zero tillage as the alternative treatment (unlike some meta-analyses that have compared reduced to conventional tillage separately e.g. van Kessel et al.^36^). When papers included multiple different tillage practices that could have been counted as a control treatment, they were further classified as conventional or reduced tillage, based on reported equipment and/or method of plowing.

#### Cover crops

Papers identified from the additional search string of “cover crop*” OR “green manure” OR “catch crop*” were included when a control treatment with no cover crop was present (e.g. bare soil when the cash crop was not growing). Experiments were included when the cover crop was grown intentionally to protect the soil and was not harvested, and residues were mechanically terminated, chemically terminated, or left as a green manure (e.g. a crop grown specifically for fertility purposes).

#### Crop rotation

Papers identified from the additional search string of “rotation” AND “continuous” were included when there was a control treatment that represented the continuous (year after year) cropping of one cash crop. The experimental treatment needed to include the same crop as well as at least one additional crop, grown in rotation (as in McDaniel et al.^23^). We included two experiments where an additional crop was grown not in rotation but as an intercrop (i.e. two plant species grown simultaneously on the same field) and one experiment that met the rotation criteria but was different in that it also included grazing in the experiment treatment but not the control (Table 1 in S1 File). In all experiments, we recorded the number of crops in rotation for analysis.

#### Perennials

Papers identified from the additional search string of “perennial” OR “agroforest*” included experiments where a perennial treatment was compared to an annual cropping system. This practice represented more significant shifts in management practices that have been the subject of fewer studies, thus we included control practices that varied slightly (for example, they included monocultures with or without conventional tillage). Treatments included perennial grasses, agroforestry and managed forestry (Table 1 in S1 File). While these treatments have differences in species and management, they share the critical feature of continuous living cover through perennials. Given the limited number of total studies, we aggregated these into a single class (as in Basche and DeLonge 2017^25^). Two of the eight experiments ultimately included in this practice also had livestock grazing as part of the treatment (compared to an annual crop system with no livestock; Table 1 in S1 File).

#### Crop and livestock

Papers identified from the additional search string of “graz*” AND “livestock” were included if there was a crop-only control and a treatment with a similar crop system that also included livestock grazing. This treatment was of interest as it is representative of one phase of integrated crop-livestock systems that has implications for diversifying cropland management. The identified studies included experiments with either annual crop or pasture-based systems, where control systems were harvested conventionally (i.e. with equipment) whereas treatments included livestock grazing and no conventional harvesting.

### Database design

Data from experiments were extracted and categorized systematically. When experiments reported measurements from several years, years were included separately. When experiments included multiple measurements of infiltration rate within a year, measurements were averaged, as has been done in other meta-analysis evaluating soil properties that may be measured on a sub-annual basis^23^. This approach, which was used for 10 studies (and 11% of the response ratios in the database), allowed us to use as much data as possible to capture the influence of the treatments on infiltration rates over a longer timeframe.

We analyzed additional variables to examine how effects of management on infiltration rate are modulated by other factors of interest^23,37,38^. These variables included soil texture (percent sand, silt, clay), climate, study location, and study length. We also analyzed additional information within select practices, including tillage descriptions (within no-till), inclusion of cover crops (within no-till), the number of crops grown in an experiment (within crop rotations), and if crop residues were removed or maintained (within cover crops). Study length was defined as the number of years a treatment was in place, as reported by the authors, and we assumed that this duration explains differences between control and treatment conditions.

We supplemented our dataset using publicly available sources to explore broader patterns that could be influencing the effectiveness of management practices. When annual precipitation was not reported, we used the Global Historical Climatology Network (GHCN)-Daily database^39^ (contains records from over 80,000 stations in 180 countries and territories). As an additional indicator of longer-term climate conditions for all study sites, we used locations to extract estimates for the aridity index, an integrated measure of temperature, precipitation and potential evapotranspiration (CGIAR-CSI Global-Aridity and Global-PET Database, resolution of 30 arc seconds^40,41^). In cases where soil textures were not reported in papers from the U.S. (which represented the largest number of studies, Table 1), we used data from the U.S. Department of Agriculture’s Web Soil Survey^42^.

### Statistical analysis

Statistical analysis was conducted by calculating response ratios, representing a comparison of control treatments to experimental treatments, as is common in meta-analysis methodology^43^. Response ratios represented the natural log of the infiltration rate measured in the experimental treatment divided by the infiltration rate measured in the control treatment (Equation 1) ^43^. A weighting factor was included in the statistical model as is suggested by Phillibert et al.^44^ and was created based on the experimental replications of each study (Equation 2)^45^. Natural log results were back transformed to a percent change to ease interpretation. Results were considered significant if the 95% confidence intervals did not cross zero.

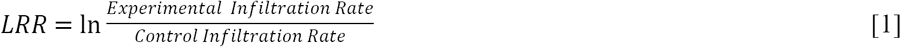

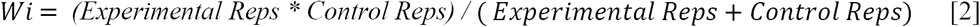

For statistical analyses, the five practices were analyzed separately because there were notable differences in experimental designs and control treatments. A linear mixed model (lme4 package in R) was used to calculate means and standard errors for the five practices. The statistical model also included a random effect of study to account for the factor of similar environments and locations in the cases where experimental designs allowed for multiple paired observations (e.g. a single study included multiple tillage practices or multiple cover crop treatments using different species)^46^. For the two practices that included the largest number of studies (no-till and cover crops) and could therefore be statistically evaluated in greater detail, additional fixed effects including mean annual precipitation, study length and soil texture, were analyzed with a similar linear mixed model^47^. Given the limited sample sizes for the other three practices (perennials, cropland grazing and crop rotations) additional fixed effects models could not be robustly applied, but figures were developed to explore trends (Figs 5-7 in S1 File). Regression coefficients were calculated to determine the effect of continuous environmental variables (Table 2). Additional details, including sample R code, are provided in the Supplementary Information.

**Table 2.** Regression coefficients (β) for continuous environmental and study variables included in the analysis. (aridity index, annual precipitation, % sand content in soils, % clay content in soils, and length of study (treatment duration) (n=number of paired comparisons per practice, bold notes p < 0.05).

| Practice | Aridity Index |  | Annual Precipitation |  | % Sand |  | % Clay |  | Study Length |  |
| --- | --- | --- | --- | --- | --- | --- | --- | --- | --- | --- |
| | $\beta$ | n | $\beta$ | n | $\beta$ | n | $\beta$ | n | $\beta$ | n |
| No-Till | 0.028 | 207 | 0.000 | 207 | 0.001 | 188 | -0.003 | 189 | 0.016 | 207 |
| Cover Crop | -0.009 | 81 | 0.000 | 81 | **0.010** | **69** | -0.015 | 72 | 0.015 | 81 |
| Crop Rotation | 1.228 | 39 | **0.001** | **39** | 0.004 | 32 | -0.008 | 38 | -0.005 | 39 |
| Perennial | 0.011 | 40 | 0.000 | 37 | 0.004 | 18 | 0.022 | 20 | -0.007 | 40 |
| Crop and livestock | 0.430 | 24 | 0.000 | 24 | 0.005 | 20 | 0.008 | 20 | 0.010 | 24 |

A sensitivity analysis was performed for each of the practices using a Jacknife technique, where individual experiments were removed from the respective databases and overall means were recalculated, to determine how sensitive effects were to individual experiments^44^. To determine if there was evidence of a preference for published studies with significant effects (publication bias), we evaluated histograms for all practices^48^.

## Results

### Database description

Through the methodical keyword-based literature search, we identified 89 studies eligible for inclusion in our database, representing 391 paired comparisons on six continents (Fig 3 and Fig 1 in the S1 File). Many experiments were in North America (31) or Asia (27), with most located in the United States (25) and India (20). More than half of the experiments and subsequent paired comparisons were no-till (207 paired comparisons from 52 studies), while the next largest practice was cover crops (81 paired comparisons from 23 studies). Sixty-three percent of the database (246/391 paired comparisons) demonstrated an increase in infiltration rate with any of the five alternative agricultural practices included in the analysis. Overall means for perennials and cover crops were significantly greater than zero (Fig 4).

**Fig 3.**
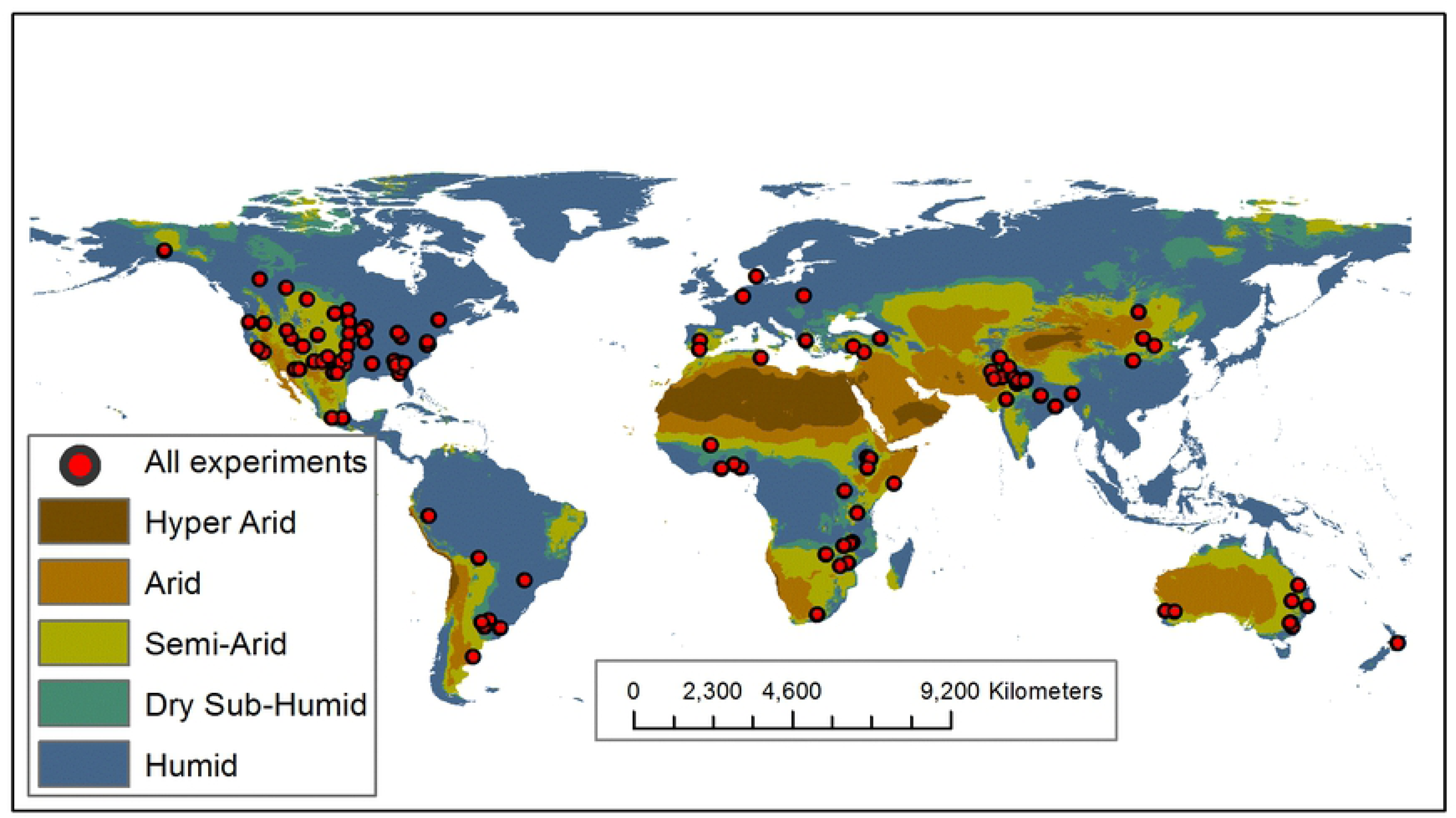
Map of experiment locations included in the analysis, with respect to their aridity regimes. Aridity regimes were determined using the aridity index, an integrated measure of temperature, precipitation and potential evapotranspiration from the CGIAR-CSI Global-Aridity and Global-PET Database^40,41^. Maps were generated with ESRI ArcGIS version 10.4 (http://www.esri.com). See Fig 1 in the S1 File for maps depicting locations for individual practices.

**Fig 4.**
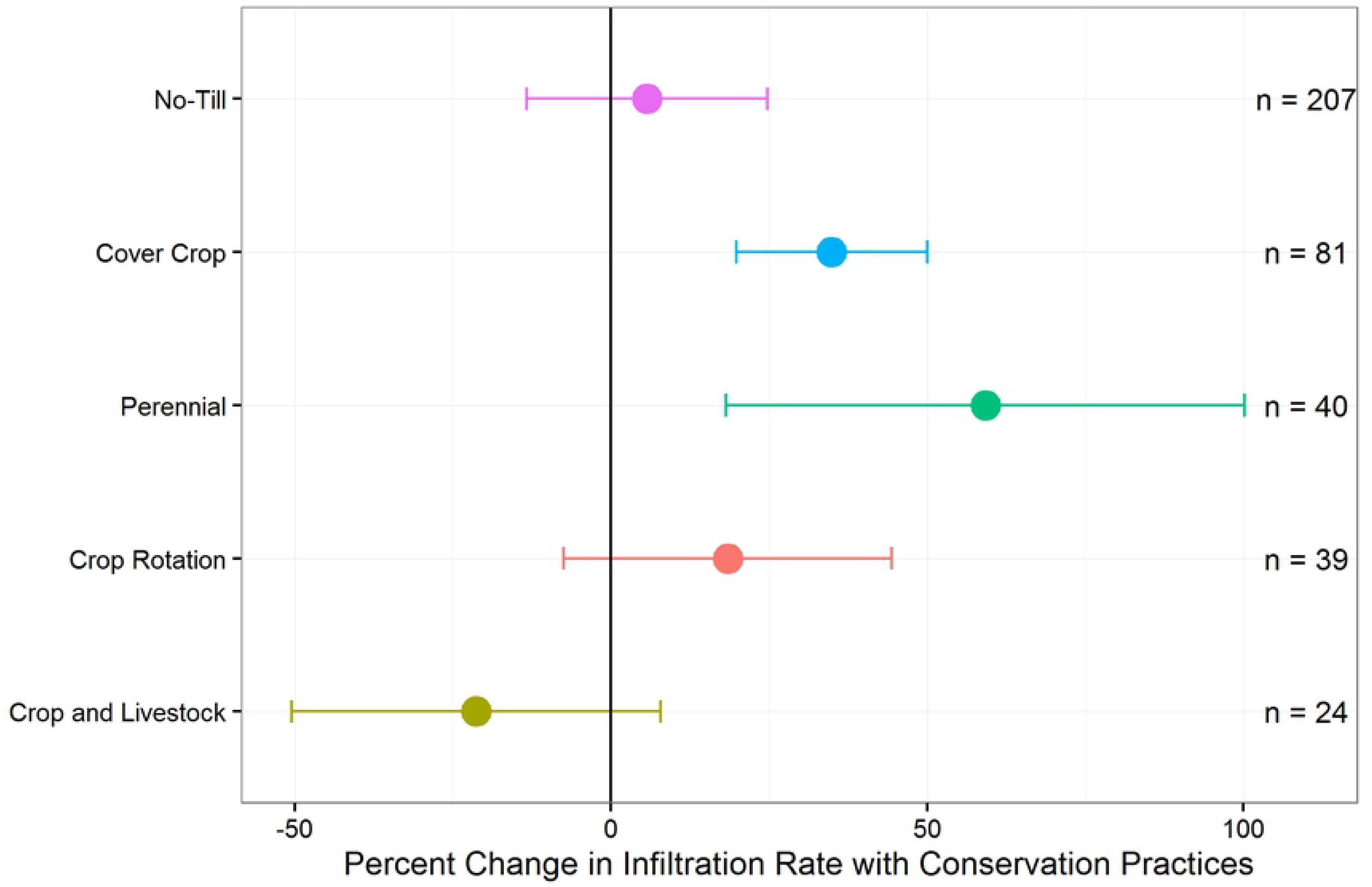
Percent change in infiltration rate with the five alternative agricultural practices included in the analysis compared to conventional controls. (mean ± 95% confidence interval, n=number of paired comparisons per practice).

**Fig 5.**
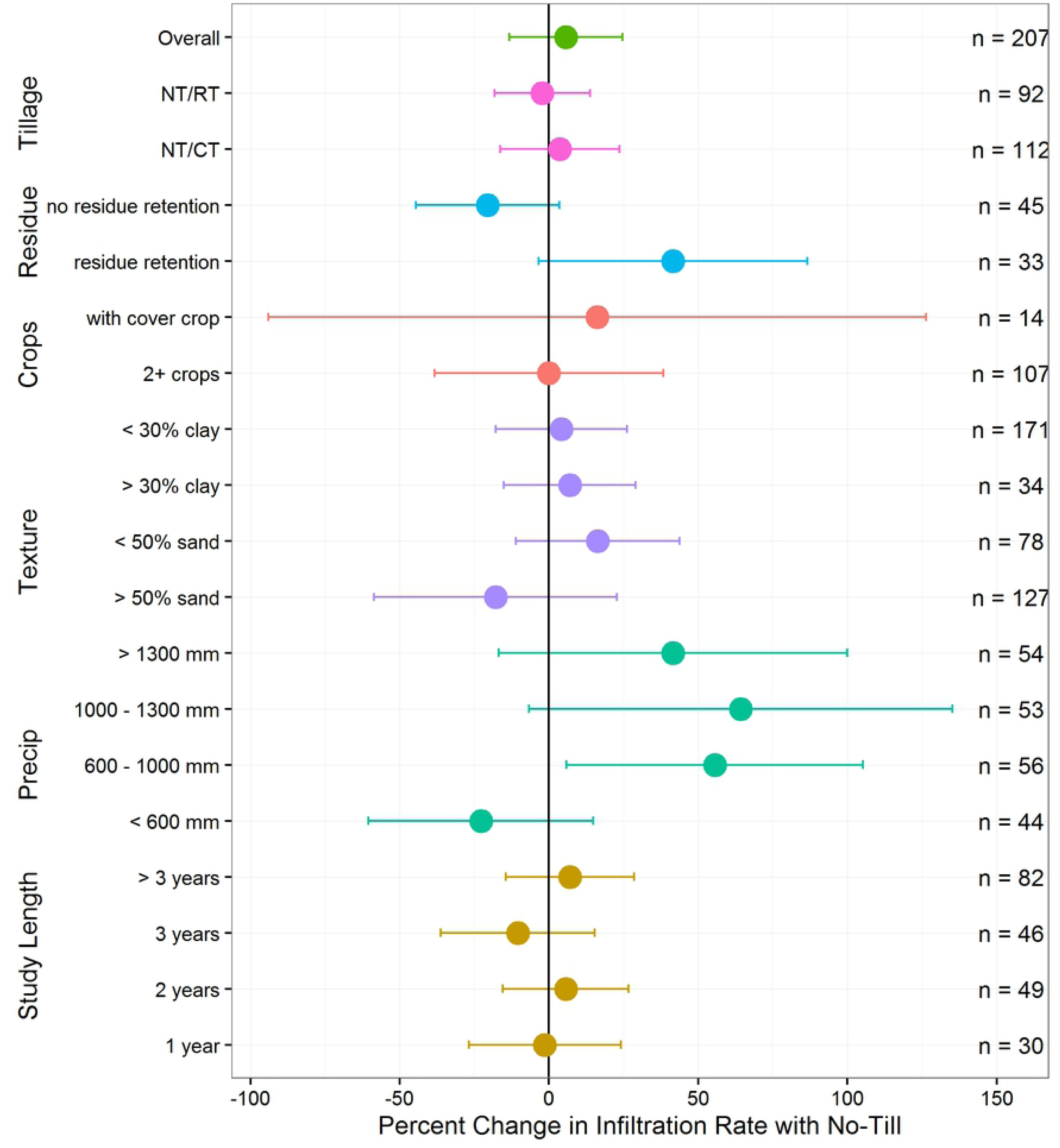
Response of infiltration rates to subsets of no-till experiments. Means and 95% confidence intervals were calculated using fixed effects for different subsets related to annual precipitation, study length, soil texture, tillage practice in controls, and crop and residue management (n=number of paired comparisons).

### No-Till

The overall mean increase in infiltration rates in no-till versus tillage comparisons was not significantly different from zero (5.7%, confidence interval −13.3 – 24.7%) (Fig 4). Also, we did not find differences between experiments comparing reduced tillage to no-till versus conventional tillage to no-till. We found the effects of no-till to be complex, revealing possible conditions and environments where no-till practices are more likely to increase infiltration rates (Fig 5). For example, in the subset of experiments reporting residue management details (11 with residue retained, 7 with residue removed), there were higher increases in infiltration rates in experiments that combined no-till with residue retention practices (41.5%, confidence interval - 3.4 – 86.6%). Only 2 of 52 experiments reported data capturing the effect of no-till plus a cover crop (compared to tillage plus a cover crop) and results were inconclusive (16.2%, confidence interval −94.0 – 126.5%). Similarly, there was no significant difference when no-till experiments included more crop diversity (in both control and experimental treatments), such as having at least two crops in rotation or double cropping (0.0%, confidence interval −18.9 – 18.8%). With respect to environmental variables, we found an effect of precipitation, with significant improvements in regions with 600 to 1000-mm annual precipitation (55.6%, confidence interval 5.8 - 105.3%) (Fig 5). There were also greater numbers of results where no-till reduced infiltration rates located in more arid environments (i.e., lower aridity indices), but the effect was not statistically significant (Table 2 and Fig 2 in the S1 File). We did not detect any clear effects of soil texture, nor did we find differences due to study length (Table 2 and Fig 3 in the S1 File).

### Cover crops

The mean increase in infiltration rates for cover crop experiments (n=81, 23 studies) was significantly above zero (34.8%, confidence interval 19.8 – 50.0%) and results demonstrated a few other important differences relative to patterns observed in no-till experiments. For example, there was a significant improvement in infiltration rates when cover crop experiments were in place for more than four years (30.0%, confidence interval 1.7 – 51.3%, representing 34 of the 71 comparisons) (Fig 6). Also, we did not detect differences when cover crop experiments were aggregated by annual rainfall or aridity index (Fig 6 and Table 2). There was evidence that the effects of cover crops on infiltration rate improvements were greater in coarsely textured soils with higher sand contents and less clay (Table 2 and Fig 3 in the S1 File). Similar to the no-till plus residue retention experiments, we found there to be a significant increase in infiltration rates when experiments combined cover crops with no-till (compared to no cover crops with no-till; 44.6%, confidence interval 11.6 – 77.5%) (Fig 6).

**Fig 6.**
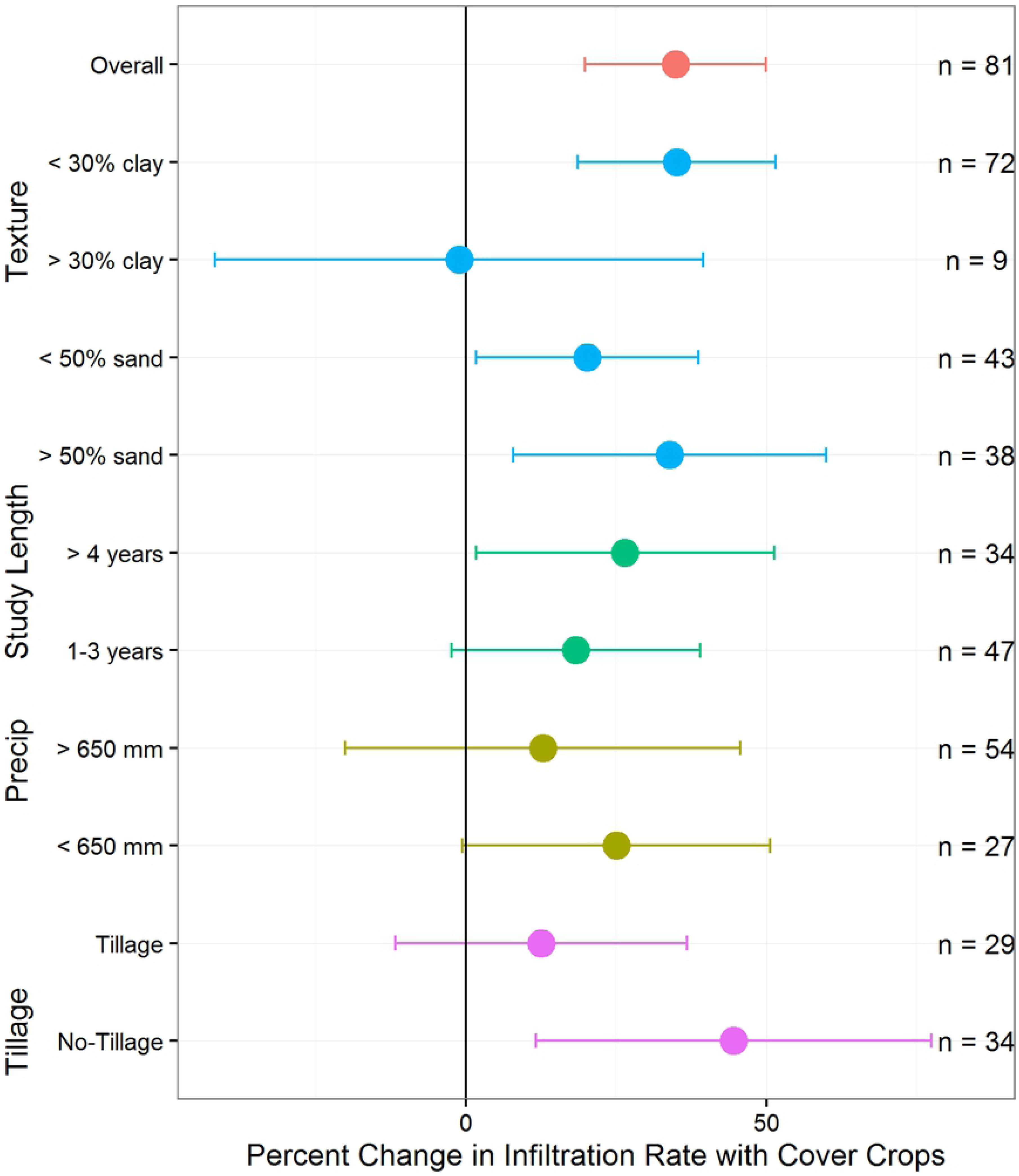
Response of infiltration rates to subsets of cover crop experiments. Means and 95% confidence intervals calculated using fixed effects for subsets related to annual precipitation, study length, soil texture, and tillage practice (n=number of paired comparisons).

### Crop rotations

Impacts of crop rotations on infiltration rates were inconsistent, with an overall mean effect that was not significantly different from zero (18.5%, confidence interval −7.4 – 44.4%, n=39 from 11 experiments) (Fig 4). Many experiments in our database compared monoculture to two crops in rotation, and only a few compared three or more crops in rotation. Further, in many experiments the control crop was monoculture maize (Fig 2 in the S1 File). The aridity index analysis revealed that most of the declines in infiltration rate among the crop rotation experiments fell within more arid regions (Table 2 and Fig 5 in the S1 File).

### Perennials

Experiments comparing perennial treatments to annual crops showed the largest improvement in infiltration rates (59.2%, confidence interval 18.2 – 100.2%, n=40 from 8 experiments) (Fig 4). These experiments included three types of perennial systems: agroforestry, perennial grasses, and managed forestry (Fig 6 in the S1 File); they were aggregated into a single group for this analysis because of the limited number of available studies (only eight total met the inclusion criteria) and because they share a key feature of continuous roots in the soil (Table 1 in the S1 file). Despite differences among and between these practices, the perennial practices showed a consistent pattern in that growing perennial rather than annual plants led to improved infiltration rates.

### Crop and livestock (cropland grazing)

Experiments that fit our criteria for crop and livestock systems were more likely to contribute to a decline in infiltration rates overall (−21.3%, confidence interval −50.4 – 7.9%, n=24 from 7 experiments) (Fig 4). However, individual studies within this practice suggested that pasture-based and diversified annual crop systems with livestock could lead to improved infiltration rates under some conditions (Fig 7 in the S1 File).

**Fig 7.**
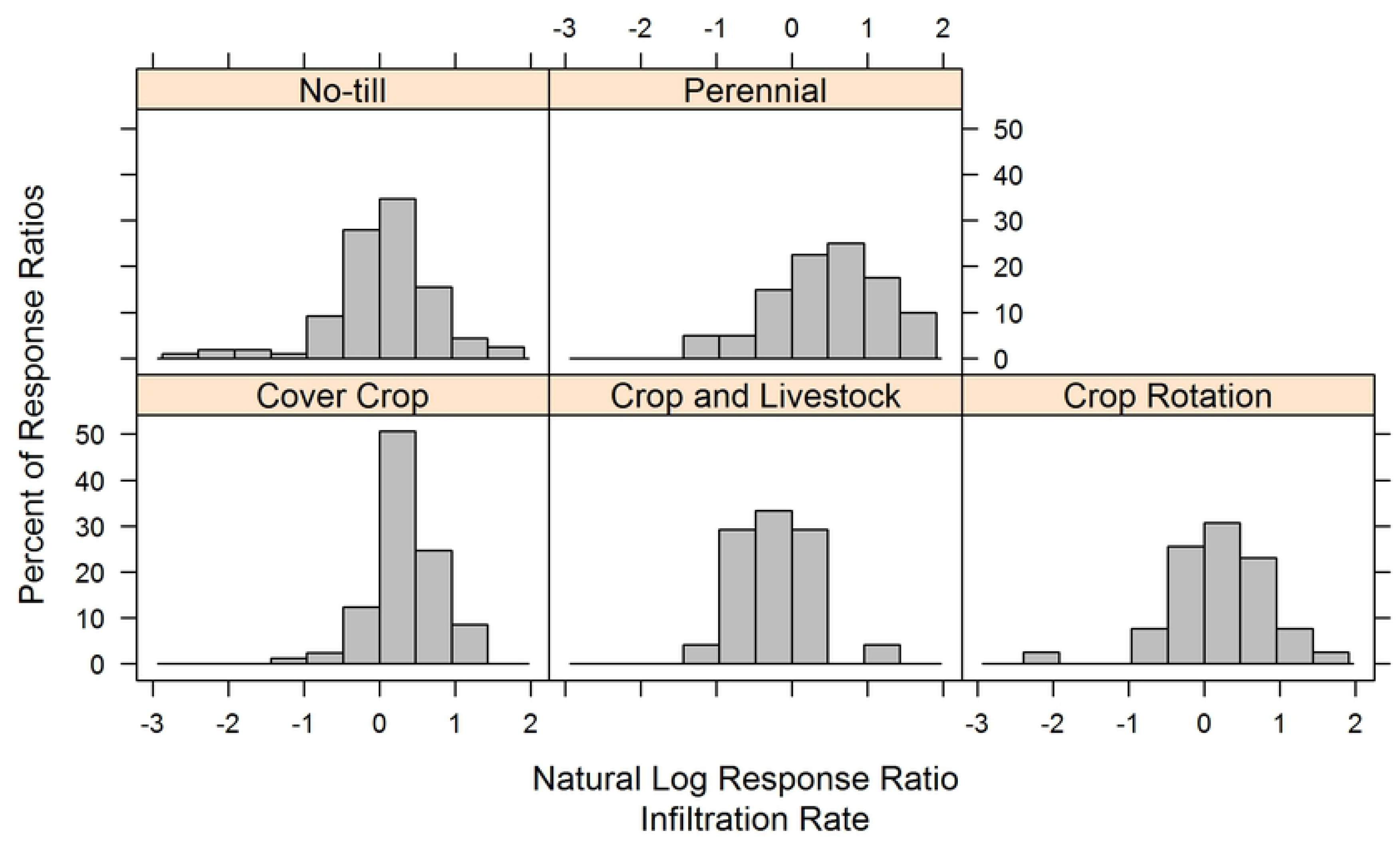
Publication bias analysis using histograms of response ratios. Histograms created using the methodology suggested by Rosenberg et al. (2000). Normal distributions indicate that publication bias was not likely a factor in study results (i.e. there was not a bias against publishing experiments that did not have significant effects).

### Publication bias and sensitivity analysis

We did not find evidence of publication bias in our overall analysis, as shown by histograms demonstrating that experimental results within each practice were not skewed toward very positive or very negative effects (Fig 7). Also, the Jacknife sensitivity analysis revealed robust results, with only minor shifts to overall means and confidence intervals when individual experiments were removed (Fig 8). Results were most robust for no-till and cover crops, which had the largest numbers of experiments. However, two practices – crop rotation and perennials – were somewhat sensitive to the removal of individual experiments. When two of the eight perennial experiments were separately removed, the 95% confidence intervals of response rates shifted to slightly cross zero (Fig 8). These experiments were the two with livestock, which suggests that in these environments the presence of livestock did not reduce infiltration^49,50^. For the crop rotation studies, the removal of one experiment^51^ led to a significantly different mean from zero.

**Fig 8.**
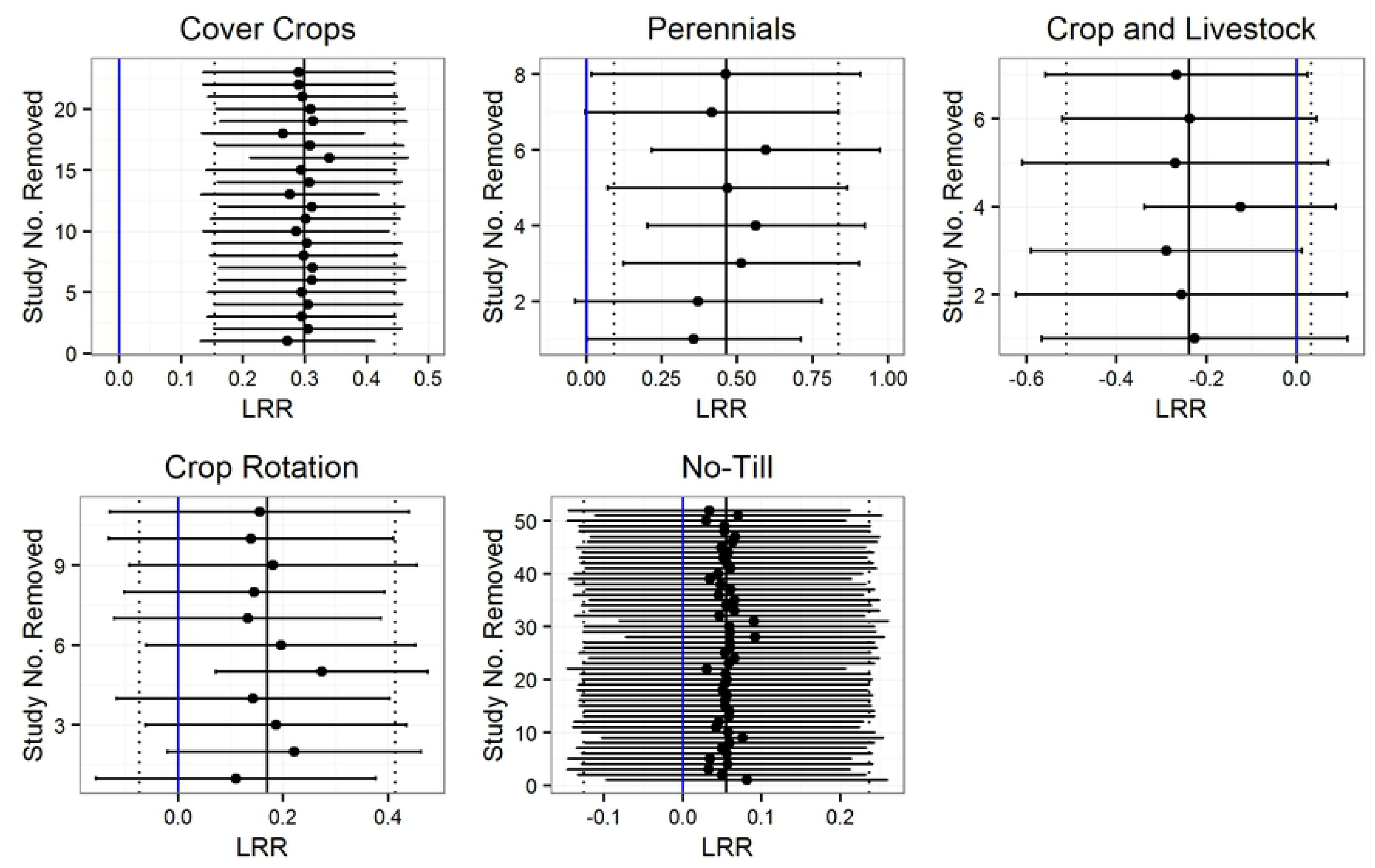
Sensitivity of results to individual studies using a Jacknife technique. Blue lines represent zero or no effect, and 95% confidence intervals that do not cross zero were considered significant. The solid black line represents the overall practice means and the dashed lines are overall 95% confidence interval before individual studies were removed to re-calculate the displayed means and confidence intervals.

## Discussion

### Alternative management impacts infiltration, likely through biological, chemical and physical processes

Overall we found that the largest infiltration rate changes were associated with practices that entail a continuous presence of roots and soil cover, suggested by the positive improvements of perennial systems compared to annual crops and cover crops compared to no cover crops, as well as the negative trend associated with the crop and livestock systems compared to crop systems only. Determining the exact processes underpinning the observed results is outside the scope of meta-analysis. However, these results point to changes in soil hydrologic function which, in turn, is known to be associated to an intertwined set of biological, chemical and physical factors. For example, physical processes associated with root growth and decomposition contribute to improved soil structure such as porosity and aggregation, which enhances water entry into the soil^52^. Recently, Basche and DeLonge^25^ found that cover crops, perennial grasses and agroforestry practices led to significant improvements in two soil hydrological properties related to water infiltration (porosity and water retained at field capacity), which could help explain the effects from those practices in this analysis. The reduced infiltration rates that we found with respect to the crop and livestock studies could be related to the removal of vegetative cover or soil compaction from grazing, although the available studies for this practice were limited^53-55^. Overall, our results suggest that management has an important contribution to infiltration rates, and that these are likely related to soil physical changes.

Given established relationships between soil carbon and soil water properties^26,27^, one factor that likely has a role in our findings is the impact of carbon accrual from the analyzed practices. For example, increases in soil carbon have been quantified by meta-analyses in response to cover crops, crop rotations, and other conservation practices^7,23,24^. Also, perennial systems typically store more soil carbon than annual croplands^56-58^. However, reviews evaluating the effect of no-till on carbon have found mixed results^22,59-62^, similar to the complex no-till findings in the present analysis. Specifically, these reviews have found that no-till can lead to carbon accrual in some instances but may also lead to no net increase in carbon but rather a redistribution of carbon closer to the soil surface^59^. Further, it has recently been demonstrated that the relationship of soil carbon to soil available water may not be as strong as indicated by prior analyses^63^.

Continuous cover of the soil combined with reduced soil disturbance is known to promote enhanced biological activity, with is also linked to physical soil structure. For example, management practices leading to a greater number of earthworms could contribute to soil aggregation and pore creation, increasing water entry^64,65^. A recent meta-analysis found that reduced tillage increased earthworm abundance and biomass by more than 100% compared to conventional inversion tillage^66^, suggesting a potential biological mechanism that may help explain the success of no-till in improving infiltration rates under some circumstances. Cover crops have also been found to increase earthworm populations and recent work finds that they also significantly increased microbial biomass as well as mycorrhizal colonization across a range of experiments.^67-69^ Similar changes in biological activity could be expected in other practices that promote crop diversity and year-round growth, such as crop rotations and perennial systems.

### Comparing the efficacy of different management practices

Our results suggest similarities and distinctions between alternative management that are in many ways corroborated with past studies that have limited their scope to a narrower range of practices. For example, the overall finding that continuous soil cover can improve infiltration rate is corroborated by prior research focused on cover crops or agroforestry. A recent meta-analysis of eight experiments in Argentina found a similar effect of cover crops on infiltration rate, where infiltration was increased by an average of 36% due to the presence of cover crops compared to no cover controls^70^. Also, Ilstedt et al.^71^ found that afforestation and agroforestry increased infiltration rates relative to annual crop systems by 100-400% across four experiments in tropical agroecosystems.

Somewhat contrary to conventional thinking around no-till, our global meta-analysis found that no-till did not consistently improve infiltration rates at this scale. In contrast to our findings, a recent qualitative review (mostly from studies within the United States, in both wetter and drier environments) found that no-till in most instances increased infiltration rates over conventional tillage^37^. Also, a review of experiments in the Argentine Pampas, a humid environment with well-drained soils, found that no-till doubled infiltration rates^38^. While our results did demonstrate a trend toward improvement, our database included very few cases where infiltration rates increased by at least a factor of two as a result of no-till, even in humid environments (16/207 paired comparisons; Table 1 in the S1 File). Also, we did not find a significant effect of no-till in the subset of no-till experiments including cover crops (Fig 5), contrary to our findings in for cover crops (where cover crops increased infiltration rates within the subset of cover crop studies with no-till, Fig 6). This inconsistency may be related to the limited number of no-till experiments reporting infiltration rates for combinations of factors, such as use of cover crops, which would have allowed more comprehensive analysis. We did, however, find that no-till experiments with residue retention were more likely to increase infiltration rates, suggesting the importance of combinations of practices to maximize benefits.

Crop rotations had an inconsistent effect on infiltration rates. We did observe a negative effect of crop rotations on infiltration rates in drier regions (Table 2; Fig 2 in the S1 File). However, the studies that met our criteria were largely from more arid regions, so the limited dataset may have inhibited analysis across a sufficiently wide range of aridity regimes in order to detect stronger overall effects. In a meta-analysis that similarly considered conventional management versus crop rotations but focused on soil carbon, McDaniel et al.^23^ found that crop rotations generally increased carbon, but that greater increases were correlated with more precipitation. Thus, the study revealed a sensitivity of crop rotation impacts to climate, potentially related to small decreases in bulk density that may have affected soil hydrologic function^23^. Together, these findings suggest a need to closely monitor the impacts of crop rotations on several soil variables, especially in drier environments. This may be especially important for this practice, as there is already great deal of variability in the crop diversity and level of complexity of crop rotation practices.

Although limited experiments fit our criteria for crop and livestock systems, the overall result suggests that careful management of these complex systems may be necessary to maintain or increase infiltration rates. While the mean change in infiltration rates was negative across all studies, individual experiments suggested that a positive effect was possible under some circumstances and management practices. For example, Masri and Ryan^72^ found infiltration rates increased when a diverse annual crop rotation included livestock as compared to when the systems included crops only. Franzluebbers et al.^73^ reported increased infiltration rates in pasture-based systems with versus without livestock, but only when a lower grazing intensity was utilized. It is also important to note that cropland grazing typically represents only one component of a diversified farming system that may have different outcomes when assessed on a larger scale.^74^

### Uncertainty surrounding measurement timing and experiment duration

One variable potentially affecting our results could be related to a sensitivity to the timing of measurements in these experiments. This sensitivity may be particularly relevant for the no-till studies. For example, immediately after a tillage event, the infiltration rate in tilled fields could increase relative to no-till because of managed decreases in bulk density^37^. An experiment included in this analysis^75^ found greater seasonal differences versus treatment differences when comparing tillage practices to no-till. Our database could not be categorized according to inter-season periods of measurement and management, as such analysis would have been complicated by inconsistent data availability and was beyond the scope of our study. As such, we were only able to evaluate overall trends based on available data and these limitations likely account for some uncertainty in our analysis.

Another related variable that could be introducing uncertainty is the lack of studies reporting effects following a wide range of treatment durations. In our analysis, we did not find experimental length to be a significant factor in our analysis across any of the practices (Table 2 and Figure 4 in the S1 File). This finding is contrary to the common convention that management practices need be in place for an extend period of time in order to demonstrate improvements to various soil properties. Instead, we found that even after a short period (as little as one year) it was possible for infiltration rates to increase relative to conventional controls. This finding could also be related to the interannual timing of measurements, as infiltration rate is a dynamic process subject to interseason and/or interannual variability. However, examining such effects was beyond the scope of this analysis, as the primary goal was to detect infiltration rate changes between different farming practices.

### Uncertainty surrounding data limitations and research gaps

Overall, our results revealed the varying relative abundance of experiments evaluating different practices; no-till experiments comprised more than half of our database, while many fewer experiments evaluated practices such as perennials or crop and livestock systems. This observation aligns with recent findings indicating that more complex agroecological research receives relatively limited research funding^76,77^. While we did find several studies for each practice, our sensitivity analysis revealed that the limited number of experiments in some led to more sensitive results. Smaller sample sizes also limited our ability to explore influences of other environmental and management factors (e.g. we were able to comprehensively evaluate the effects of precipitation and soil texture only for no-till and cover crop practices).

Additional levels of analysis that also consider the combined and synergistic effects of multiple management practices would also be valuable. For example, it would be interesting to compare the combined effects of no-till, cover crops, and crop rotations (typically combined in conservation agriculture systems) as compared to conventional agricultural systems. However, such analysis was beyond the scope of this study and would be challenging given the very limited number of experiments that combine practices and report results in a sufficiently similar way to directly compare controls and treatments. More complex, well-replicated, and long-term studies would be needed to enable a similar meta-analysis to the present study, but with this broader scope.

In general, a lack of detail on environmental and management factors was another important gap in our analysis. Gerstner et al.^78^ and Eagle et al.^79^ proposed criteria that field experiments should include to increase their utility for meta-analyses or synthesis reports, in the fields of agronomy and ecology. These criteria include environmental features, such as soil and climate characteristics, as well as reporting complete factorial results from experiments.

## Conclusions

The overall trend quantified by this analysis is the potential for improvements to infiltration rates with various alternative agricultural management practices, with the greatest benefits observed in response to introducing perennials or cover crops. Our findings suggest the importance of the presence of continuous living plant roots and the positive physical soil transformations that accrue as a result. We found that no-till practices did not consistently increase infiltration rates but were more likely to do so in more humid environments or when combined with residue retention. Another important finding is that some practices have been substantially less studied than others, particularly ones that show some of the greatest promise for facilitating water infiltration and therefore mitigating the effects of extreme weather that are expected to grow more frequent with climate change.

## Acknowledgments

We would like to thank Joy McNally for providing support in the literature search, Jasmin Gonzalez and Oliver Edelson for support in database development, and Dan Kane for his feedback on prior versions of this work.

## Supporting Information

The Supporting Information (S1 File) document includes descriptions of all experiments, maps of experimental locations for the five different practices, model selection and R code, figures depicting the continuous variables included in the analysis, and additional analysis of the perennial, crop rotation and crop and livestock systems.

## Data Availability Statement

IR_meta.csv contains the database utilized in this analysis.

## Author Contributions

A.B. and M.D. designed the analysis. A.B. performed the literature search. A.B. and M.D. conducted the analysis and wrote the manuscript.

## Additional Information

Competing interests: The authors declare that they have no competing interests.

